# Parameter inference for enzyme and temperature constrained genome-scale models

**DOI:** 10.1101/2022.07.05.498798

**Authors:** Jakob Peder Pettersen, Eivind Almaas

**Affiliations:** Department of Biotechnology and Food Science, NTNU- Norwegian University of Science and Technology, Trondheim, Norway; K.G. Jebsen Center for Genetic Epidemiology, Department of Public Health and General Practice, NTNU - Norwegian University of Science and Technology, Trondheim, Norway

## Abstract

The metabolism of all living organisms is dependent on temperature, and therefore, having a good method to predict temperature effects at a system level is of importance. A recently developed Bayesian computational framework for enzyme and temperature constrained genome-scale models (etcGEM) predicts the temperature dependence of an organism’s metabolic network from thermodynamic properties of the metabolic enzymes, markedly expanding the scope and applicability of constraint-based metabolic modelling.

Here, we show that the Bayesian calculation method for inferring parameters for an etcGEM is unstable and unable to estimate the posterior distribution. The Bayesian calculation method assumes that the posterior distribution is unimodal, and thus fails due to the multimodality of the problem. To remedy this problem, we developed an evolutionary algorithm which is able to obtain a diversity of solutions in this multimodal parameter space.

We quantified the phenotypic consequences on six metabolic network signature reactions of the different parameter solutions resulting from use of the evolutionary algorithm. While two of these reactions showed little phenotypic variation between the solutions, the remainder displayed huge variation in flux-carrying capacity. This result indicates that the model is underdetermined given current experimental data and that more data is required to narrow down the model predictions. Finally, we made improvements to the software to reduce the running time of the parameter set evaluations by a factor of 8.5, allowing for obtaining results faster and with less computational resources.

## Introduction

Temperature is a key effector of life, which is partially due to the consequence that temperature has on catalytic properties of enzymes. For a long time, it has been known that enzymatic reactions slow down at low temperatures, whereas high temperatures destroy the enzymes, rendering them non-functional. In recent research^1, 2^, it has also been acknowledged that enzymes have lower catalytic rates at high temperatures due to changes in heat capacity. The effect of temperature on the behaviour of microorganisms as a whole is evident. Freezing food stops spoilage by inactivating microorganisms, whereas cooking kills them. In between these temperature extremes, there are observable effects which can be utilized commercially. One example of this is yeast production of aroma compounds, which has been shown to depend on temperature^3, 4^, a finding with potentially great impact on wine and beer brewing.

Until recently, no attempt has been made to computationally explain the temperature dependence of microorganismal phenotypes by propagating the temperature dependence of metabolic enzymes to the entire metabolic network of an organism. However, Gang Li *et al*.^5^ came up with an extension of an enzyme-constrained genome-scale metabolic model (ecGEM) which can capture the temperature dependence of metabolism. This model is thus called an *enzyme and temperature constrained GEM* (etcGEM). As most other models, the one by Li *et al*.^5^ is based on sets of assumptions and parameters. In particular, this model is based on model ecYeast7.6^6^ (*Saccharomyces cerevisiae* strain S288C) and contains 764 metabolic enzymes and 2,292 parameters associated with enzymatic activity. For each of the enzymes, the following parameters must be determined: (1) *T_m_*, the melting temperature; (2) *T_opt_*, the temperature optimum; and (3) 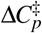, the change of heat capacity from the ground state to the transitional state.

Given these parameters, it was demonstrated how an enzyme’s maximum catalytic rate *k_cat_*(*T*) at a certain temperature *T* could be estimated^5^. These temperature-dependent maximal catalytic rates were then fed into the enzyme-constrained genome-scale model^6, 7^, and the metabolic flux rates were predicted using Flux Balance Analysis (FBA)^8, 9^. Furthermore, Li and coworkers^5^ used a Bayesian approach to infer the enzymatic parameters mentioned above from sets of training data: (1) Maximal growth rates in aerobic batch cultivations^10^; (2) Maximal growth rates in anaerobic batch cultivations^11^; (3) Chemostat cultivations which include measurements of exchange fluxes of carbon dioxide, ethanol and glucose^12^.

In the training data, the experimentally determined exchange fluxes were recorded for a range of temperatures, thus generating a set of growth scenarios. The performance of a parameter set was assessed by predicting the flux rates in these growth scenarios and comparing these fluxes rates with the experimental results. Hence, the *R*^2^ score between the experimental and modelled fluxes were used to assess the model’s goodness of fit.

For defining the Bayesian model, prior parameters have to be chosen for the enzymes. Li *et al*.^5^ did this through a custom heuristic which was partially based on measured temperature optima and denaturation temperatures for enzymes. By training the model with the Sequential Monte Carlo based Approximate Bayesian calculation method, an estimate of the posterior distribution of parameter sets was found^5^. However, Li *et al*. did not systematically investigate the stability of this calculation method, nor whether it suffered from identifiability issues. Thus, it is unclear how metabolic flux results from the etcGEM can be interpreted.

In this paper, we investigated the stability of the Approximate Bayesian calculation method algorithm by choosing multiple different random seeds over different priors. We found that the Bayesian calculation method is inherently numerically unstable, and thus, is unable to provide reliable results given its model assumptions. To rectify this problem, we implemented an evolutionary search algorithm that is not built upon any assumption of structure of the underlying data. Finally, we improved the execution time of the software package by almost a factor 10, making it feasible to execute on smaller-scale computational infrastructures. However, for the available data there is an identifiability problem, in which solutions that equally match the experimental data still differ in terms of fluxes through key metabolic reactions. We believe that the evolutionary algorithm will resolve the identifiability problem if more experimental data, in particular data regarding internal fluxes, are included.

## Results

### Improvements to the running time of the algorithm

Running the Bayesian calculation method once for 500 iterations with the chosen hyperparameters consumed approximately 17,000 CPU hours (approximately corresponding to two weeks on a 48 core computer) using the implementation from Li *et al*. Profiling showed that the particle evaluation procedure (see Methods) was the performance bottleneck, and excess time consumption was caused by COBRApy’s^13^ internal routines to modify metabolic models prior to solving them. Hence, preparing the models for optimization consumed far more time than the optimization proper. For this reason, we modified the implementation to use the ReFramed package (https://github.com/cdanielmachado/reframed) for handling the genome-scale model. We benchmarked the two versions on a computer running Intel Core i7-8565U using a single core (Table 1). With our code improvements, the performance was boosted by factor of 8.5. As a consequence, the results of the Bayesian calculation method could be obtained the day after starting it when running on a compute server. Still, only about 20% of the particle evaluation time was spent on optimization, so improvements within the ReFramed package has the potential for increasing performance even more.

**Table 1.** Comparing benchmark results for a full evaluation of a particle using all three conditions (aerobic, anaerobic and chemostat). The numbers were averaged over 10 iterations.

| Implementation | Overall time [s] | Percentage time in optimization [%] |
| --- | --- | --- |
| Original version from Li *et al.* | 106 | 0.837 |
| Updated version with ReFramed | 13.4 | 19.7 |

### Assessing the stability of the Bayesian calculation method

In order to investigate the stability of the Bayesian calculation method for stochastic effects, we ran the Bayesian calculation method with two different random seeds on the three training datasets. These runs of the Bayesian calculation method are referred to as Bayesian simulation 1 and 2. Also, in addition to using the priors selected by Ref.^5^, we created three randomized priors by permutation (see Methods for details) and repeated the process of assessing stability given these priors. Thus in total, we ran the Bayesian calculation method eight times, yielding eight different populations of estimated posterior distributions. The permuted priors yielded approximately the same rate of fitness convergence as the unpermuted priors (Figure 1 **A** and **B** and Supplementary Figure S1). Between simulation 1 and 2 for the same priors, the differences were negligible.

**Figure 1.**
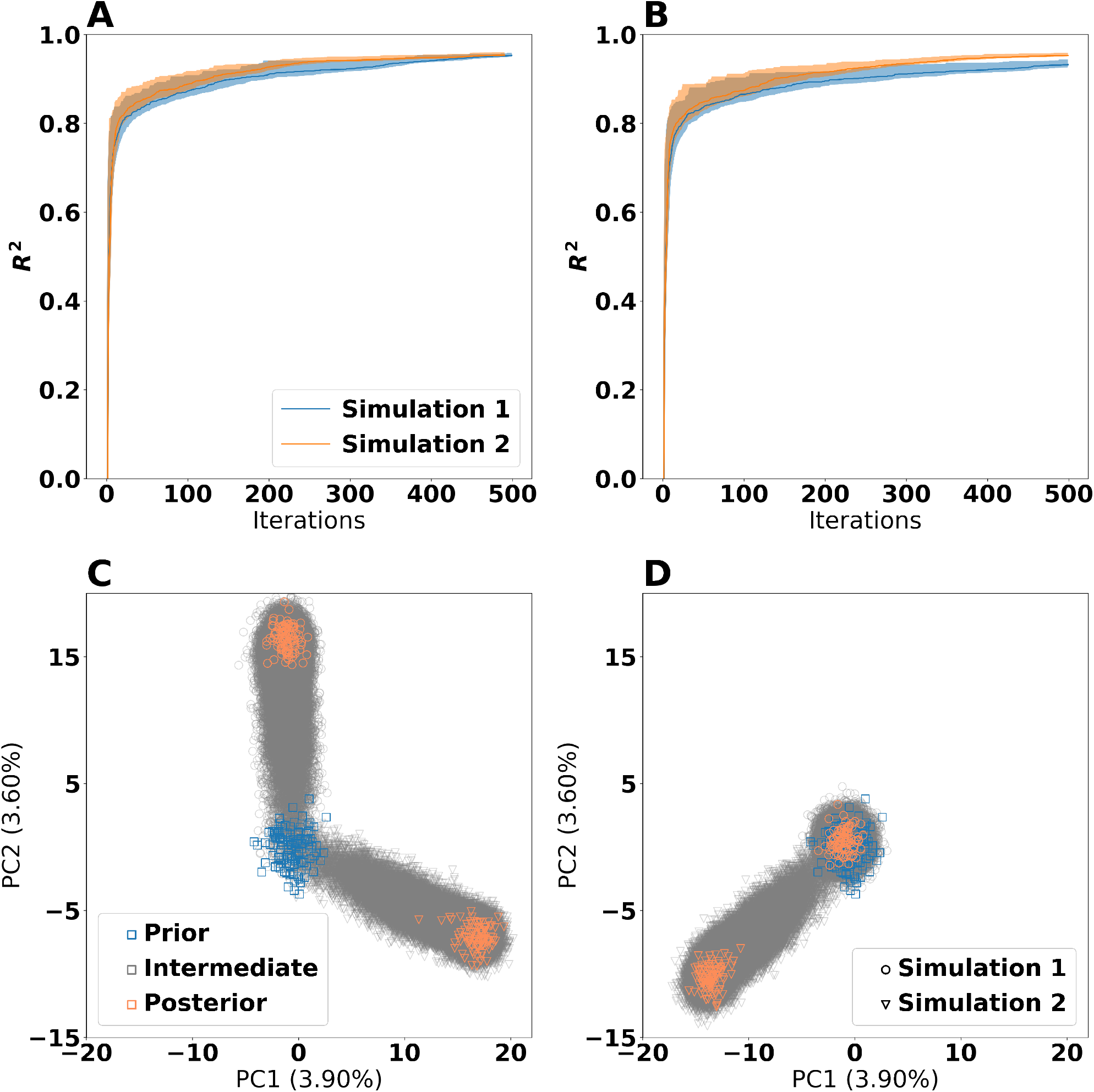
Comparison of the effect of prior and random seed on the Bayesian calculation method. Panel **A** and **B** show the training *R*^2^ values for the unpermuted priors and permuted prior 1, respectively. An *R*^2^ value of 1 corresponds to exact correspondence between the training data and the model predictions. The shaded regions indicate the 5^th^ and 95^th^ percentiles, whereas the solid lines indicate the median (50^th^ percentile). Panel **C** and **D** show Principal Component Analysis (PCA) plots of the parameter sets from the unpermuted priors and permuted prior 1, respectively. Each point is a candidate parameter set. The prior points are the ones which served as a starting point for the calculation method, the estimated posterior points are the ones which had *R*^2^ > 0.9, whereas all other points are intermediate points stemming from the simulations. The axes are identical for both panels and use the same ordination, making the panels directly comparable.

Having observed that all priors do indeed result in parameter sets with high fitness, we next investigated whether these solutions were similar. For this, we created a Principal Component Analysis^14^ (PCA) plot of the parameter sets obtained under estimation (Figure 1 **C** and **D** with a more complete overview in Supplementary Figure S2). The estimated posterior distributions, defined as the collection of particles having *R*^2^ > 0.9, were different for every simulation even though the convergence properties were similar. This means that the Bayesian calculation method is unstable for all four priors and that there are identifiability issues causing the calculation method to converge at different locations in the parameter space.

We also discovered that the calculation of *R*^2^ values for the chemostat dataset suffered from numerical instabilities unrelated to the Bayesian calculation method. For the same particle and software version, the Gurobi solver could sometimes judge the model infeasible given the parameters and sometimes it could find a feasible solution. However, given that a solution was found, the results were consistent up to expected numeric accuracy. We therefore suspected that the inherent numerical instability in calculation of the *R*^2^ value for chemostat data in turn had made the Bayesian calculation method unstable. Given this concern, we also ran the Bayesian calculation method without the chemostat data. We only used the priors suggested by Li *et al*. and ran four different simulations with differing random seed. The results for this setup (Supplementary Figure S3 and S4) were similar to the simulations including the chemostat dataset. Hence, the Bayesian calculation method was unstable also when withholding the chemostat dataset.

### Assessing enzyme-level stability of the Bayesian calculation method

To systematically study the phenotypic behaviour of particles in the estimated posterior distributions, we performed Flux Variability Analysis (FVA)^15^, see Methods for more details. While it is impossible to lock down a specific flux distribution due to an infinite number of alternative optimal solutions, FVA uncovers the flux bounds of each individual reaction capable of supporting optimal metabolic behaviour, in this case maximizing the growth rate given the model parameters. We decided to focus the investigation of FVA results on six reactions that have important biochemical roles in the metabolic network:

- Pyruvate dehydrogenase: A key reaction in connecting glycolysis to the TCA cycle and fatty acid synthesis
- Fructose-bisphosphate aldolase: An intermediate reaction in glycolysis
- Ferrocytochrome-c:oxygen oxidoreductase: The oxygen consuming reaction in the respiratory electron transport chain
- Phosphoserine phosphatase: Reaction producing the amino acid serine from intermediates in the glycolysis
- Shikimate kinase: Intermediate reaction in synthesis of folate and aromatic amino acids (phenylalanine, tyrosine and tryptophan)
- Growth: The growth (biomass) reaction is included for comparison with the other reactions. The calculated growth should ideally be identical to the experimental ones, but some deviations occurred because the posterior particles did not in general provide a perfect fit to the data.

We chose to focus on three simulations; Bayesian simulation 1 and 2 with original priors and Bayesian simulation 1 with permuted prior set 1 (Figure 2). First, we observe that the flux through shikimate kinase had a narrow flux range and was highly coupled with growth. This reaction is a part of the shikimate pathway for producing folate and aromatic amino acids. We suspect that the resulting compounds have no functionality in the model except for being part of the biomass reaction. As no alternative pathways for producing these compounds exist, the flux through the shikimate kinase reaction is thus locked at a certain fraction of the growth rate. For the other reactions, there exists more variability among the solutions. Fructose-bisphosphate aldolase and Ferrocytochrome-c:oxygen oxidoreductase are for some particles used extensively, but in other cases not at all, still giving rise to approximately the same growth rates regardless. This means that the metabolic model uses alternative pathways depending on the choice of enzyme thermodynamic parameters. The fluxes for Pyruvate dehydrogenase and Phosphoserine phosphatase generally follow the trends of the growth curve, as for shikimate kinase. However, there are outliers deviating from this pattern, again most likely due to availability of alternative pathways.

**Figure 2.**
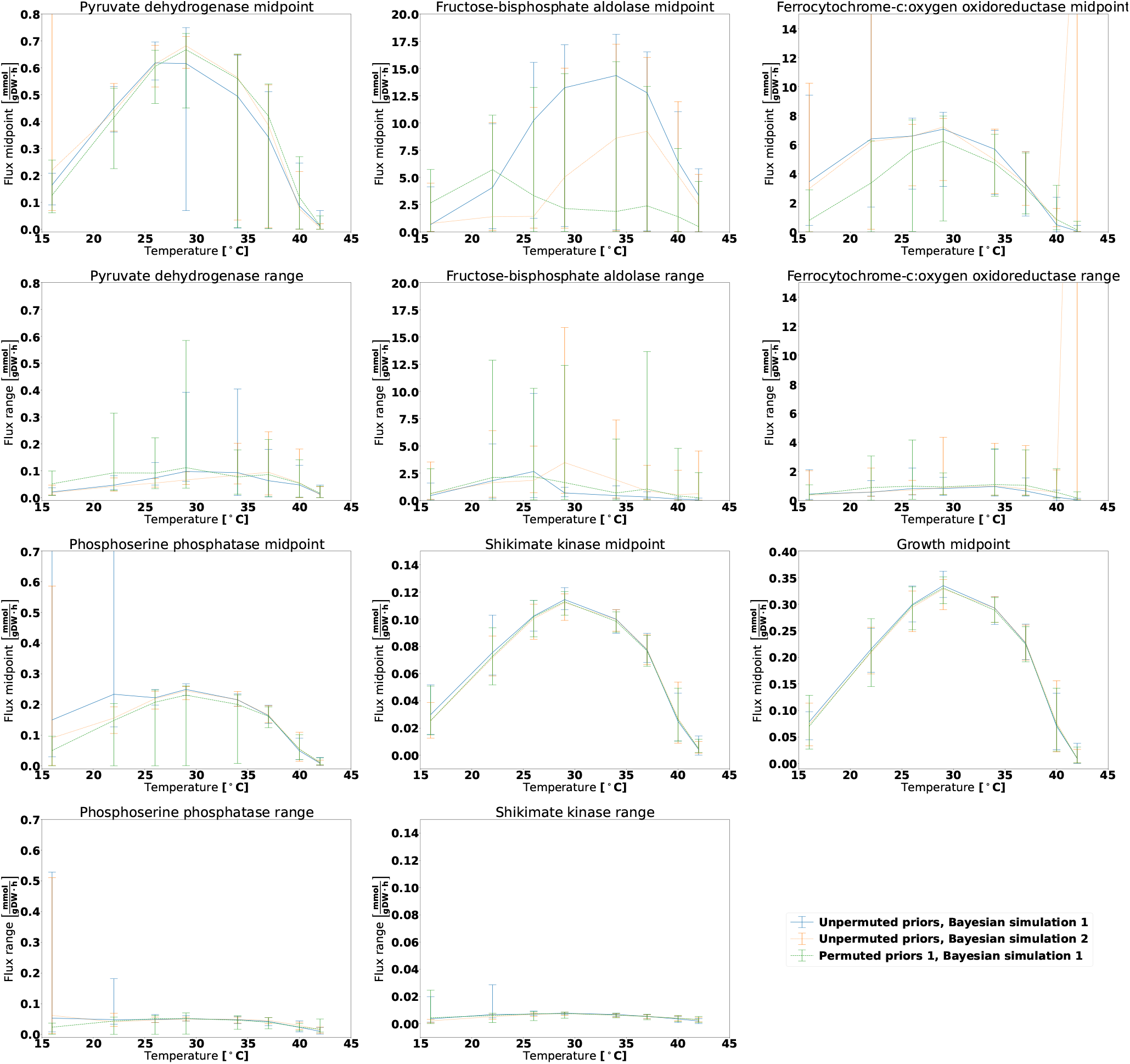
FVA analysis for the aerobic dataset for six reactions and varying temperature when using estimated posterior distributions obtained for the Bayesian calculation method. The midpoint panels show the FVA flux midpoint, this is: The average of the maximum and minimum attainable flux given the optimization objective. The range panels show the absolute difference between the maximum and minimum flux. The lines denote the mean midpoint or range value, whereas the error bars span from the lowest to the highest observed value. The growth reaction is included for reference, and it will always display an FVA range of zero as it is the optimization target.

The results for the anaerobic dataset (Supplementary Figure S5) were similar, except for the fact that there was no flux through the Ferrocytochrome-c:oxygen oxidoreductase reaction as there was *no* oxygen available to be consumed. We also ran FVA on the results omitting the chemostat dataset when running the Bayesian calculation method (Supplementary Figure S6 and S7). These results also showed large variability within simulation results and across simulations.

### A bimodal toy example

Given the observation that the Bayesian calculation method returned different parameter sets with high fitness, we suspected that the fitness landscape of the temperature parameters was multimodal. Hence, we decided to test the Bayesian calculation method on a toy problem with two parameters to infer; *x* and *y*.

We defined the fitness function as:

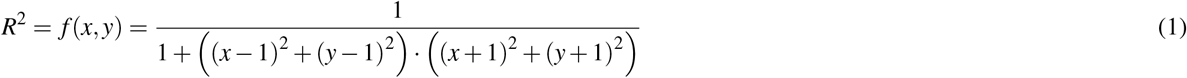

This fitness function assumes values from 0 to 1 where *R*^2^ = 1 is only attained in the global maxima (*x, y*) = (1, 1) and (*x, y*) = (−1, −1). The function does not have any additional extrema, but it has a saddle point at (*x, y*) = (0, 0).

As our priors, we assumed that *x* and y were independent and identically normally distributed with mean zero and standard deviation 0.2. This is:

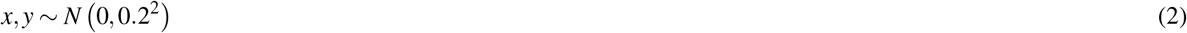

Due to the symmetry of this problem, the true posterior distribution is equally centred around the two optima and we would therefore expect the Bayesian calculation method to replicate this symmetric distribution. We ran the Bayesian calculation method on this toy problem with a population size of 32 over 200 iterations with four replicates having different random seeds. When plotting the final generation of particles (Figure 3 **A**), we realized that the Bayesian calculation method clustered all of its points in a very small space close to one of the optima. Which of these two optima this was, varied based on the random seed, but the same simulation never yielded points near both of the optima. In addition, only one of the four simulations actually reached an optimum, whereas the three other simulations suffered from genetic bottlenecking, meaning that the variability in the population of particles disappeared and caused premature convergence^16^.

**Figure 3.**
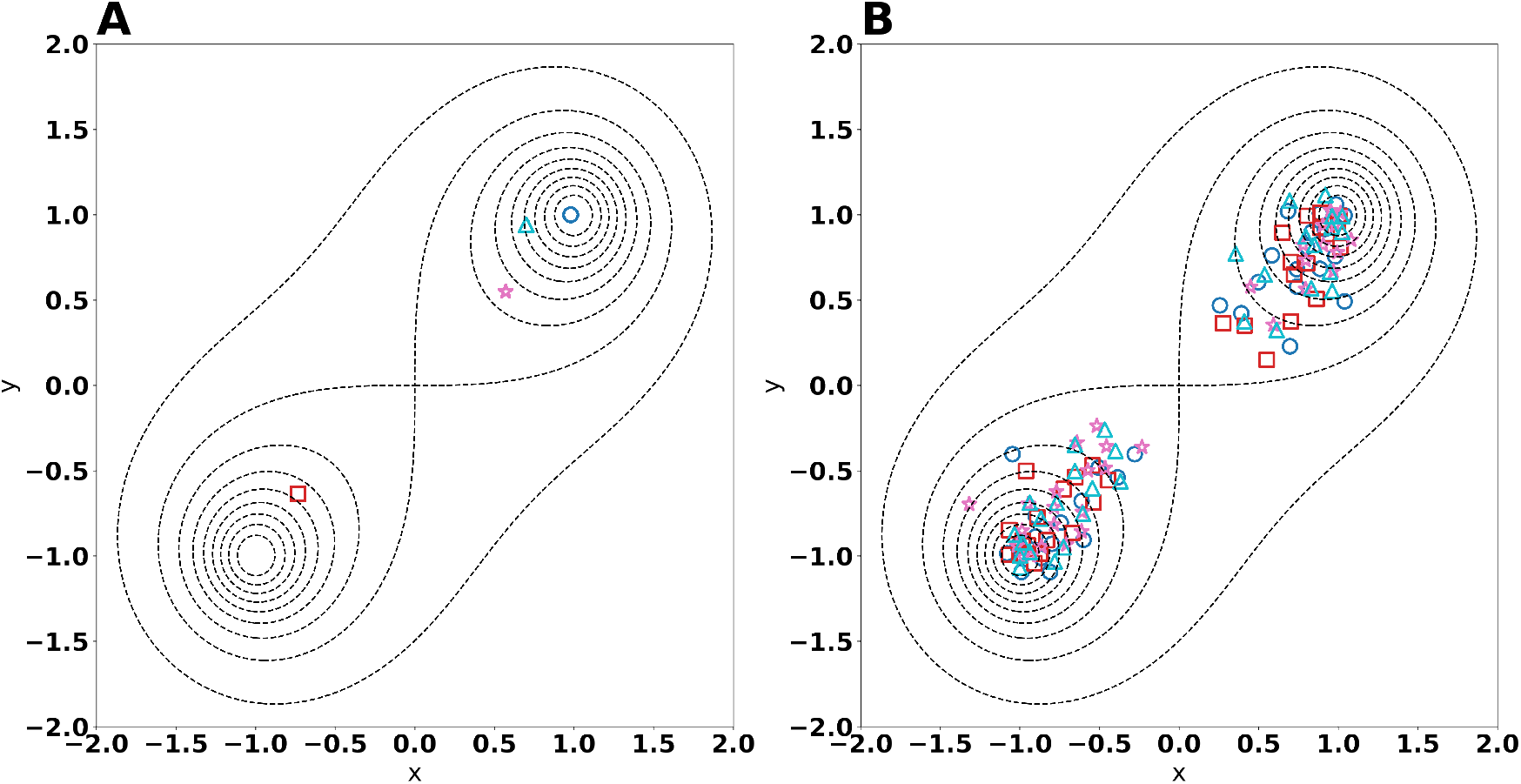
Results of parameter inference on the toy example. Panel **A** and **B** show the final generation of particles for the Bayesian calculation method and evolutionary algorithm, respectively. Each point represents an individual in the final population. Each shape and colour (each shape is associated to exactly one colour) represents one of the four replicate simulation. The global fitness optima of the problem are the points (−1, −1) and (1, 1), and are marked by dotted contours. For Panel **A**, the particles from each simulation are so close that they are visually indistinguishable and therefore appear as a single point.

### Evolutionary algorithm

Given the instability of Bayesian calculation method and its inability to cope with multimodal fitness landscapes, we constructed an evolutionary algorithm^17–20^ as an alternative for inferring parameters. More specifically, we used a variation of CrowdingDE^21^ which is designed to find alternative optima in a multimodal distribution^22^ (see Methods for details). Our choice of an evolutionary method was motivated by how an evolutionary algorithm searches the parameter space and its ability to combine existing solutions to create improved solutions^23^. CrowdingDE has two major hyperparameters, the scaling factor *F* and the crossover probability *CF* which both determine how crossover between individuals is done.

For testing the performance of the evolutionary algorithm, we used the previously mentioned toy example with the same priors. As for the Bayesian calculation method, the population size was set to 32 and four replicate simulations were run for 200 generations. The scaling factor was set to 0.5, the crossover probability was 0.5 and 16 new children were born per generation. (Figure 3 **B**) As opposed to the Bayesian calculation method, the evolutionary algorithm diverged into two subpopulations closing on the two optima, each consisting of approximately half the individuals. This shows that the chosen evolutionary algorithm is able to find multiple optima during the same simulation. Note however, that the two subpopulations had a considerable variability after 200 generation and thus did not suffer from genetic bottlenecking.

Encouraged by the results from the toy example, we applied the evolutionary algorithm on the problem of finding enzyme parameters. We used the same prior as suggested by Li *et al*. and chose to discard the chemostat dataset for these simulations due to its associated instability. The population size was set to 128, the children born per generation was 64 and the simulation were run for 1000 generations. We varied the hyperparameters scaling factor *F* and crossover probability *CF*. Simulations were conducted in replicate, meaning that for any combination of scaling factor and crossover probability, two simulations were executed with differing random seeds.

All simulations produced particles with *R*^2^ > 0.9 by 1000 generations (Supplementary Figure S8). However, there were considerable variability in fitness among the particles in each simulation and only in two of the simulations (the ones with *F* = 0.5 and *CR* = 0.99), the median population fitness exceeded *R*^2^ = 0.9. Still, having a large span of fitness values inside the same simulation is not a major disadvantage per se as one can selectively pick the individuals with high *R*^2^. At the same time, having a large variability among the solutions is preferable to avoid genetic bottlenecking. The choice of hyperparameters also affected the rate of convergence. From what we can assess, *F* = 0.5 and *CR* = 0.99 yielded the best effect in this case (Figure 4 **A**), and we proceeded with the results from this hyperparameter combination. However, this does not necessarily mean that better choices for the control hyperparameter do not exist, nor does it mean that these hyperparameter values are appropriate given different experimental data sets^24^.

**Figure 4.**
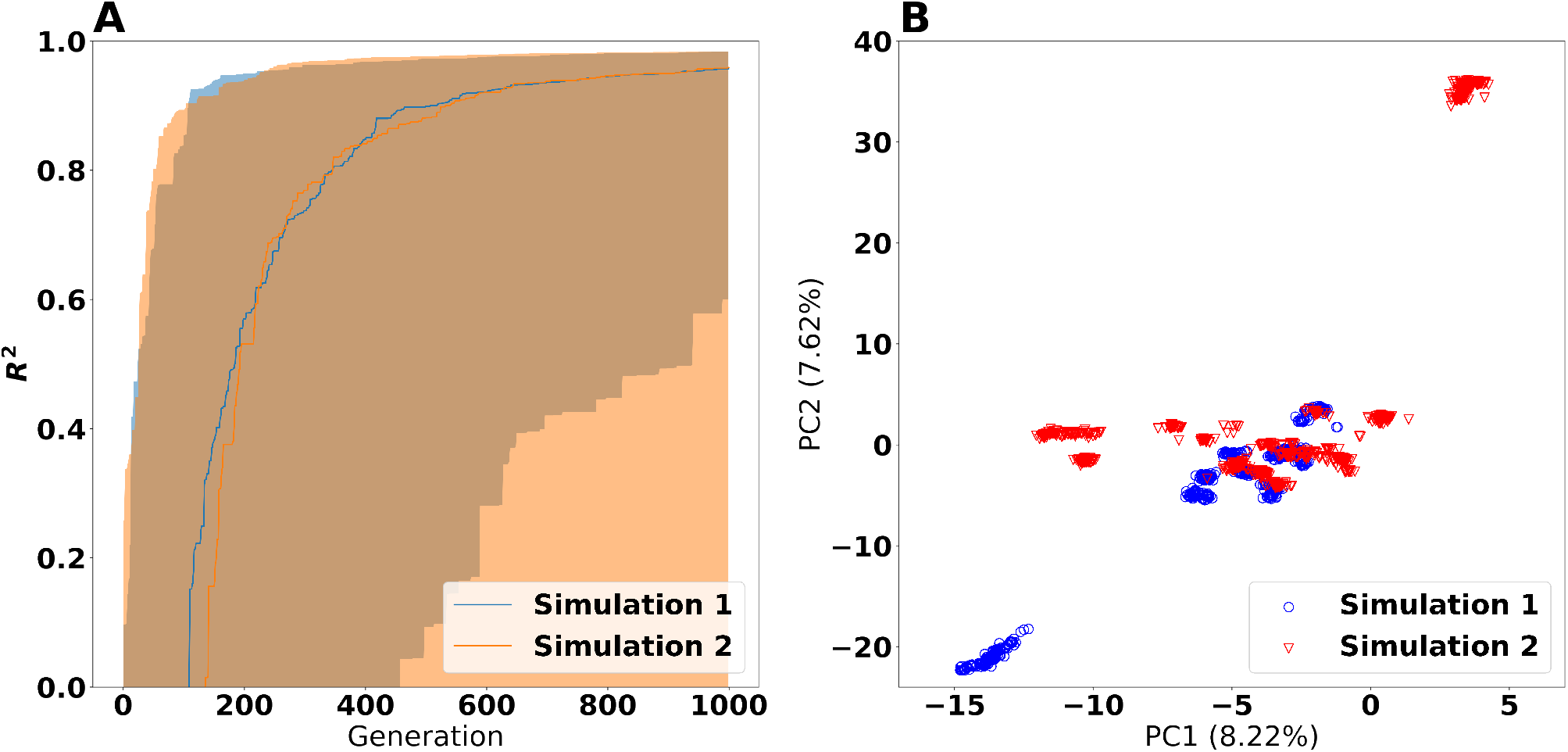
Results from the evolutionary algorithm with *F* = 0.5 and *CR* = 0.99. Panel **A** shows the training *R*^2^ values. An *R*^2^ value of 1 corresponds to exact correspondence between the training data and the model predictions. The shaded regions indicate the 5^th^ and 95^th^ percentiles, whereas the solid lines indicate the median (50^th^ percentile). Panel **B** shows the Principal Component Analysis (PCA) plot of the particles having *R*^2^ > 0.98.

We further extracted the particles from these two simulations having *R*^2^ > 0.98 and created a PCA ordination (Figure 4 **B**). From this ordination, we observed that the particles ended up in distinct clusters which we believe to be local optima of the fitness function, similar to the situation in Figure 3 **B**. Furthermore, hierarchical clusters (Supplementary Figure S9) display the particles in these discrete optima. Each simulation found a number of these optima, but the same optimum was not found by both of the simulations. Still, there is no evident distinction between the populations from the two simulations as a whole. This observation is most likely due to the fact that there are so many optima that it is not feasible for the evolutionary algorithm to find all of them.

FVA analysis of the same particles (Figure 5 and Supplementary Figure S10) revealed that the populations of particles from the two simulations did not show any large systematic differences. However, *within* the same population of solutions, there were considerable variation and outliers. This points to usage of alternative pathways which the experimental data could not lock down based on the growth rates alone. The results for Ferrocytochrome-c:oxygen oxidoreductase under aerobic condition illustrate this case; at temperatures below 37°C there were moderate levels of agreement between the different particles. However, at higher temperatures, the coupling disappeared, meaning that alternative pathways could take over and attain approximately the same fitness.

**Figure 5.**
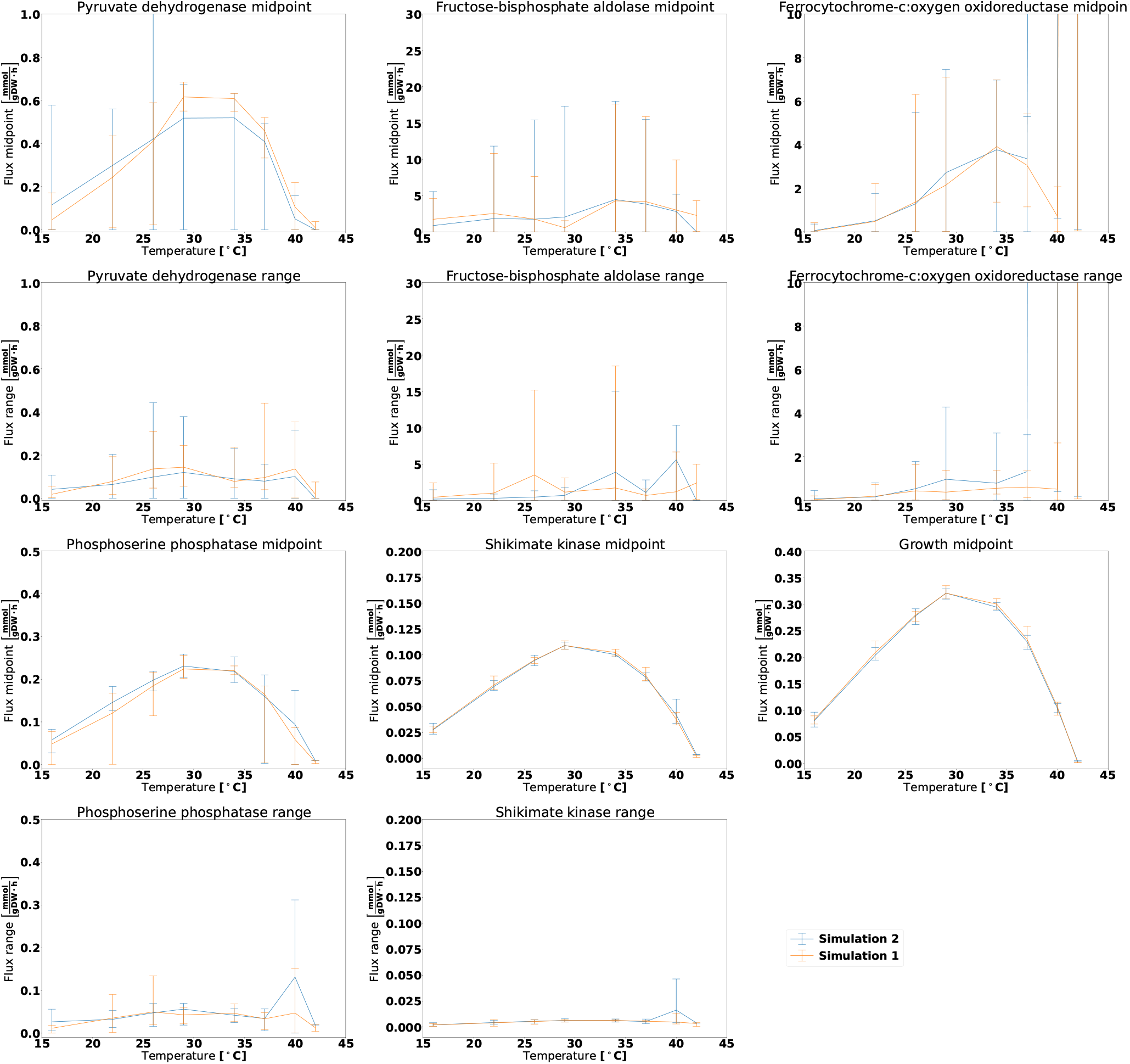
FVA analysis on the results from the evolutionary algorithm under aerobic conditions. The particles selected for this analysis stem from the two simulations with *F* =0.5 and *CR* = 0.99, considering only the particles with *R*^2^ > 0.98. The midpoint panels show the FVA flux midpoint, ie. the average of the maximum and minimum attainable flux given the optimization objective. The range panels show the absolute difference between the maximum and minimum flux. The lines denote the mean midpoint or range value, whereas the error bars span from the lowest to the highest observed value. The growth reaction is included for reference, and it will always display an FVA range of zero as it is the optimization target.

## Discussion

We observed that computing *R*^2^ values for the chemostat dataset resulted in numerical instability, while this problem was not present for the other two datasets. This is likely due to the the three-stage procedure of first locking the growth rate of the model to the dilution rate, then minimizing glucose uptake and setting it as a constraint for the model, before finally minimizing the protein usage and then reporting the fluxes. Even if the resulting problem is mathematically solvable, the sharp constraints still cause problems for the Gurobi LP solver, which for the same particle sometimes managed to find a feasible solution to the problem, and sometimes not. Potentially, this challenge could be mitigated by reformulating the optimization problem to obtain a growth rate and glucose uptake rate as close as possible (but not necessarily equal) to the target values^25^. Also, Gurobi supports directly setting lexicographic objectives solved in sequence, an approach which hopefully does not possess the aforementioned problem.

Our results point out that the outcome of the Bayesian calculation method is unstable and its result indeed depends on the choice of random seed. It is important to note that this instability has nothing to do with the instability of *R*^2^ computations for the chemostat dataset. As illustrated by the toy example, even simple bimodal fitness functions can cause the Bayesian simulations to suffer from genetic bottlenecking and failing to estimate the posterior distribution. Given the strong indications that the fitness landscape for the thermodynamic enzyme parameters is multimodal, our results imply that the Bayesian calculation method failed to converge to the true theoretical posterior distribution. This is a serious problem, as the usual statistical interpretations of the Bayesian approach will lead to erroneous conclusions if applied to the results.

Each Bayesian simulation converged to a point cloud with no apparent higher-dimensional structure. This observation makes sense, considering that the Bayesian calculation method creates new particles by sampling each parameter independently from a normal distribution where the mean and standard deviation is determined by the past generation. Thus, the estimated posterior points are likely to cluster in a high-dimensional cloud where the density of each parameter is normally distributed and the spatial density is the product of the marginal densities of the parameters. Hence, the Bayesian calculation method assumes an unimodal posterior distribution and will therefore fail when applied to a problem with a multimodal posterior distribution. The failure of the chosen Bayesian calculation method does not imply that a Bayesian approach for etcGEMs necessarily is bad, but it would require considerable refinements to the Bayesian calculation method to work with multimodal posterior distributions^26, 27^.

The evolutionary algorithm produced results which were more robust to the choice of random seed. Simulations which were run for the same combination of hyperparameters had similar development of *R*^2^ values. Although the choice of hyperparameters had large effects on the rate of convergence, we were only able to evaluate a small number of hyperparameters due to the large computational burden associated with running the evolutionary algorithm on the problem in question. We may therefore have missed out on more favourable hyperparameter combinations. Consequently, we see potential in using a self-adaptive Differential Evolution algorithm which does not need predetermining niching hyperparameters^28^.

For our preferred hyperparameter set *F* = 0.5 and *CR* = 0.99, we obtained a large variety of solutions and fitness values. The particles with the highest fitness values were dispersed among many distinct optima in the fitness landscape. Due to the high number of optima being present, the evolutionary algorithm was unable to find all of them in a single simulation. Yet, unlike the Bayesian calculation method, the evolutionary algorithm did not appear to have any spatial bias with respect to where these optima were localized in the parameter space.

The great variety of different particles of the evolutionary algorithm is not a weakness per se, but rather an indication of the desired feature of exploring the parameter space and avoiding genetic bottlenecking. Still, the results reveal that the identifiability of the parameter inference problem is poor. As revealed by FVA, there exists large variability between the particles with high fitness with respect to the internal fluxes. Hence, the choice of metabolic pathways for yeast appear to be sensitive to the thermodynamic properties of the enzymes even if the growth rate, metabolic network model topology, and external conditions were kept the same. This phenotypic sensitivity to thermodynamic properties of the metabolic enzymes may be a possible explanation for cellular metabolic heterogeneity observed in yeast cultures^29, 30^. Still, we believe that the main source for the lack of identifiability is a result of the external measurements being insufficient to account for the inner workings of the yeast cell. In this respect, we believe that measurements of proteomics^31^ and fluxomics^32^ will help narrow down the solution space and provide more accurate predictions of metabolic behaviour. For instance, if we got direct measurement of a reaction showing high variability between particles with our present data, such as Fructose-bisphosphate aldolase, we would be able to rule out the particles not satisfying the measured fluxes of this reaction.

The application of such refined approaches strongly suggests that a strategy for including different kinds of data is needed. Heckmann *et al*. inferred apparent *k_cat_* values from large-scale proteomics and metabolomics data on a genome-scale level using gene knock-out strains and machine learning. As a result, more accurate prediction of *in vivo* fluxes were obtained compared to using *k_cat_* values measured *in vitro*^33, 34^. We believe that this strategy can be adapted to the current etcGEM framework and provide efficient integration of different kinds of data, thus allowing narrowing down the solution space and at least partially alleviate the problem of identifiability.

With our software improvements, running the inference algorithm is much faster than with the original implementation, yet the problem is still so computationally expensive that workstation or server-grade hardware is required. This computational burden is likely to increase when incorporating more experimental data to calibrate the parameters. Therefore, systematic improvements should be implemented in the framework to minimize unnecessary overhead in order to hold the computational burden at a manageable level.

## Methods

### Evaluation method for particles

A parameter set, referred to as a particle, is a collection of the three parameters *T_m_, T_opt_* and 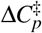 for the 764 metabolic enzymes of the ecYeast7.6 model. The goodness of fit for a particle was evaluated by the framework created by Li *et al*.^5^. In brief, this evaluation procedure acted as a black box taking a particle as input and returning an overall *R*^2^ value. In the assessment, each particle was matched against experimental data from three experiments with *Saccharomyces cerevisiae*. These were:

- The aerobic dataset^10^ measuring growth rates of yeast at 8 temperatures between 16°C and 42 °C under aerobic batch fermentations.
- The anaerobic dataset^11^ measuring growth rates of yeast at 13 temperatures between 5 °C and 40 °C under anaerobic batch fermentation. However, in our particle assessment, only 8 of the temperatures were used.
- The chemostat dataset^12^ measuring exchange fluxes of carbon dioxide, ethanol and glucose at 6 temperatures between 30°C and 38.5 °C in aerobic chemostats.

The parameters were used to adjust the effective catalytic rate *k_cat_*(*T*) (which includes denaturation) for each enzyme in the model and at each temperature *T*. In addition, the model’s non-growth associated ATP maintenance (NGAM) was also adjusted according to the temperature. Further details are published in Li *et al*.^5^.

For the aerobic and anaerobic datasets, the model’s temperature-dependent parameters were tuned, and fluxes were predicted by Flux Balance Analysis (FBA) calculations at the specific temperatures, and the biomass (growth) function was set as the objective. For each of the aerobic and anaerobic datasets, the growth rates predicted by the model were compared with the experimental growth rates, and an *R*^2^ value was reported, yielding 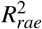 and 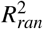.

For the chemostat dataset, the procedure was somewhat different. Here, the temperature-dependent parameters were tuned (as with the other datasets) before the growth rate of the model was locked to the dilution rate of the chemostat. Thereafter, the model was optimized for minimum glucose uptake, and this uptake flux value was set as a constraint for the model. Finally, the model was optimized for minimum protein pool usage, and exchange fluxes of ethanol, carbon dioxide, and glucose were recorded. Once the three fluxes for all temperatures in the chemostat dataset were recorded, these values were compared to the experimental ones, and 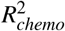 was determined.

The overall *R*^2^ for all datasets was calculated as the arithmetic mean of 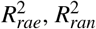, and 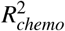. This value was then returned as the final result of the evaluation procedure. The higher the *R*^2^ value, the higher correspondence exists between the modelled solutions and the experimental results, where *R*^2^ = 1 corresponds to the highest achievable fitness.

We optimized the evaluation procedure to use the ReFramed package instead of COBRApy^13^ in order to reduce overhead related to modifying models. However, the results generated by our modified particle evaluation approach should be identical to the results generated by the original code by Li *et al*. for all simulations.

### Approximate Bayesian calculation method

The framework and code for the Sequential Monte Carlo based Approximate Bayesian calculation method was taken directly from Li *et al*.^5^, and we used the same hyperparameters as in the original publication. For seeding the calculation method, priors were needed for the values of *T_opt_*, *T_m_* and 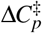 for each enzyme. We used the same priors as Li *et al.*. These priors considered the distribution of each parameter *x_i_* to be normally distributed

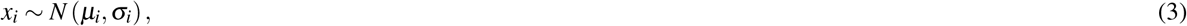

and the marginal distribution of each parameter was independent. Some simulations were also run with permuted priors. This meant that the labels of the enzymes were randomly shuffled and each enzyme thus got the values of *T_opt_*, *T_m_* and 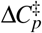 of another enzyme before this new prior was used to seed the Bayesian calculation method. The computations were run for 500 iterations. The population size at the end of each iteration was 100. We generated 128 new particles for each iteration and evaluated them according to the description in the previous section. The new particles were generated by computing the mean and standard deviation for each parameter of the particle population and sampling new parameters from a normal distribution with the aforementioned mean and standard deviation. When creating new particles, we nevertheless made sure that they obeyed the constraint *T_m_* > *T_opt_* > 0K. If this constraint was violated, the parameter was resampled. Selection of the particles was implemented through truncation selection, meaning that the 100 best particles from the previous iteration were passed to the next iteration while the rest were discarded.

### Evolutionary algorithm

The evolutionary algorithm used in this paper for fitting enzyme parameters is based on the existing CrowdingDE^21^ algorithm and was written from the ground up. Individuals in the evolutionary process were the parameter set particles discussed earlier. The population size for each iteration (generation) was set to 128. The initial 128 individuals of the population were generated by sampling from the same priors as those used by Li *et al*. The algorithm was run for 1000 generations. Each generation consisted of the following steps carried out in sequence:

- **Generation of children:** At the beginning of each generation, 64 children were created as a weighted difference of parent individuals. For each child, three parents *P*_1_, *P*_2_ and *P*_3_ were selected at random from the population without replacement, ensuring that the parents were unique. We refer to *P*_1_ as the primary parent and *P*_2_, *P*_3_ as the crossover parents. At first, each parameter for the child was initialized to the corresponding value for the primary parent, this is:

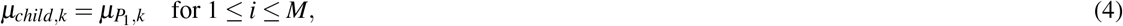

where *M* denotes the total number of enzyme parameters. For crossover, a random integer *i* ∈ [1, M] was uniformly drawn. A counter variable *j* was thereafter initiated to zero and the following procedure was repeated: A random number *r* ∈ (0, 1) was uniformly drawn. If *j* ≥ *M* or *r* > *CR*, where CR is referred to as the *crossover probability*, crossover was cancelled and the algorithm advanced to commence the generation of the next child. Otherwise, crossover was performed on parameter *k* = *i* + *j* mod *M*:

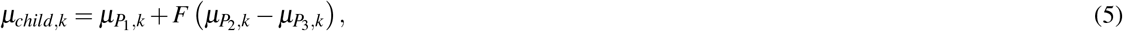

*and j* was incremented by one and the procedure was repeated for the next enzyme parameter. Updates to enzyme parameters which violated the constraints *T_m_* > *T_opt_* > 0K were reverted.
- **Evaluation of children:** Each child particle was evaluated through the same procedure as mentioned above and their respective *R*^2^ values were reported. In our case, we withheld the chemostat dataset and therefore only averaged the 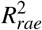 and 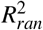 values.
- **Replacement:** For each child *a* generated in the same generation, the normalized parameter-space distance from the child to the individuals in the current population was calculated:

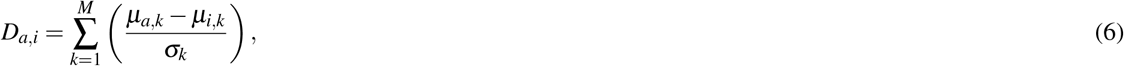

where *μ_a,k_* and *μ_i,k_* are values for the enzyme parameter *k* for the two individuals, *M* is the total number of enzyme parameters and *σ_k_* is the empirical standard deviation of enzyme parameter *k* in the population at the end of the previous generation. The closest individual *b* to the child *a* was then chosen as:

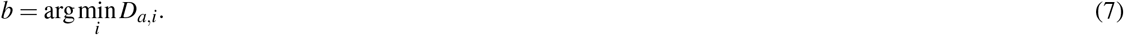 If the individual (the child) *a* had a higher fitness than b, i.e. 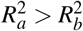, then *b* was discarded from the population and replaced with *a*. Otherwise, *b* was kept in the population and the child *a* was discarded.

The two tuning parameters *F* (scaling factor) and *CF* (crossover probability) were tested with values *F*∈ {0.5, 1.0} and CF∈ {0.9, 0.99, 0.999}. For each combination of these parameters, two replicate simulations were conducted with different random seeds.

### Ordinations

We used Principal Component Analyses (PCA) from scikit-learn (version 1.0)^35^ to create ordinations of particles. The values for each parameter were standardized, subtracting the mean and dividing by the population standard deviation before ordination. The means and standard deviations were computed across all particles present in the ordination in question. As a result, the presented ordinations in Figure 1 and Supplementary Figure S2 are comparable across the panels in the same figure. For the Bayesian calculation method (Figure 1 and Supplementary Figure S4), all points generated during the simulations were included to make the ordinations. However, for the evolutionary algorithm (Figure 4), only the points attaining *R*^2^ > 0.98 from the two simulations with *F* = 0.5 and *CR* = 0.99 were included in the ordination.

### Flux variability analysis

Flux variability analysis (FVA)^15^ was conducted on particles having *R*^2^ > 0.9 (for the Bayesian calculation method) or *R*^2^ > 0.98 (for the evolutionary algorithm) in order to ensure that these particles had high fitness. From each simulation, 20 particles were sampled randomly from the particles satisfying the aforementioned thresholds. For each sampled particle, FVA was run for the aerobic and anaerobic datasets across the same temperatures used for determining *R*^2^ in the parameter fitting process. For the chemostat dataset, the numeric instability was too large to produce reliable results, and the chemostat dataset was thus discarded for the analyses. The temperature and the parameters of the particles were first used to fix the effective *k_cat_* values. Subsequently, the metabolic model was optimized for maximal growth, and the lower bound of the growth reaction was locked to the obtained growth rate. Thereafter, maximum and minimum fluxes through each reaction in the model were found given the constraints. Instead of using the maximum and minimum fluxes directly, we converted them into flux midpoint (average of maximum and minimum) and flux ranges (absolute difference between maximum and minimum). Results with missing or infinite values were removed.

All flux ranges and flux midpoints with the same combination of simulation, reaction, dataset, and temperature were aggregated to give the mean, minimum and maximum values. Usually, there were 20 such values for each combination, as 20 particles were sampled for each of these combinations. However, this number could be smaller due to removal of missing and infinite values.

### Hierarchical clustering

Agglomerative hierarchical clustering^36^ was conducted on the particles from the evolutionary algorithm with *F* = 0.5 and *CR* = 0.99 that satisfied *R*^2^ > 0.98, as for PCA and FVA. We standardized each parameter value by subtracting the mean value and divided by the standard deviation among the selected particles and also calculated pairwise Euclidean distances. Hierarchical clustering was conducted by single (minimum distance) linkage in order to put emphasis on the detection of discontinuities between clusters of particles. The results were presented in a dendrogram showing the particles as leafs. The branches of the dendrogram were coloured according to which simulation the downstream branches corresponded to. Branches containing particles from both simulations were left uncoloured (gray).

## Acknowledgements

Thanks to Gang Li for troubleshooting technical problems with the simulations. Thanks to Ingelin Steinsland for consulting on Bayesian statistics. J.P.P. and E.A. are grateful to ERA CoBioTech project CoolWine and Norwegian Research Council grant 283862 for funding.

## Author contributions statement

E.A. and J.P.P. conceived the project. J.P.P did the analysis, created visualisations and wrote the first draft of the paper. All authors contributed to and accepted the final version of the paper.

## Additional information

### Availability of source code and data

The source code for the analysis and visualisation is available at GitHub (https://github.com/AlmaasLab/BayesianGEM). Also, an archived copy of the repository is available at Figshare (https://doi.org/10.6084/m9.figshare.21436623). Simulation of genome-scale models has been carried out with ReFramed (https://github.com/cdanielmachado/reframed) using Gurobi (Gurobi Optimization, LLC) as solver. The data required for reproducing the figures in the main article and in the Supplementary material is available as a Figshare repository (https://doi.org/10.6084/m9.figshare.21436668).

