## Supplementary Materials for "Parameter inference for enzyme and temperature constrained genome-scale models"

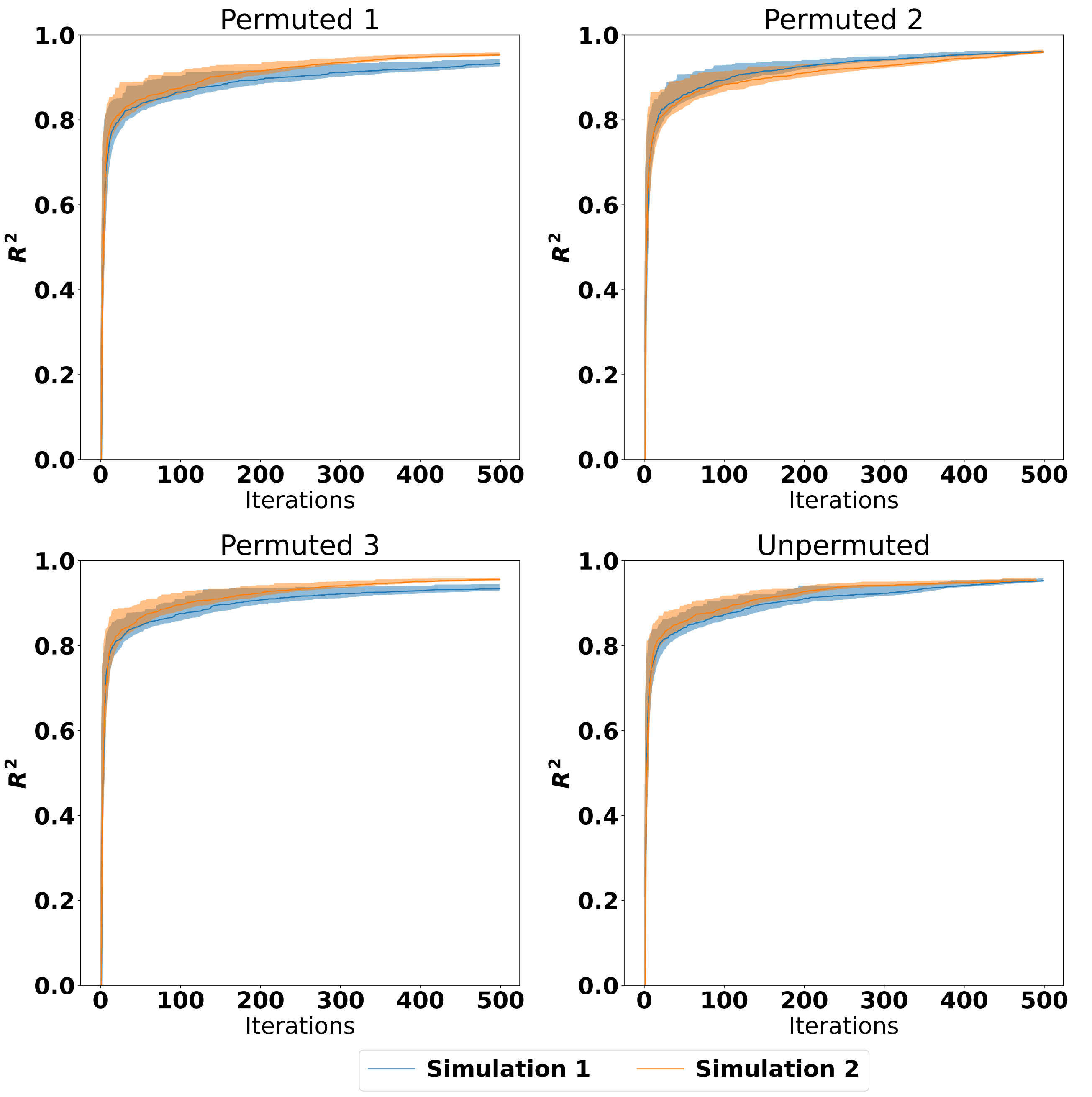


**Figure S1.** Training $R^2$ values for the Bayesian calculation method from each the four different priors. Each panel contains the results of both simulation 1 and 2. An $R^2$ value of 1 corresponds to exact correspondence between the training data and the model predictions. The shaded regions indicate the 5$^{th}$ and 95$^{th}$ percentiles, whereas the solid lines indicate the median (50$^{th}$ percentile).

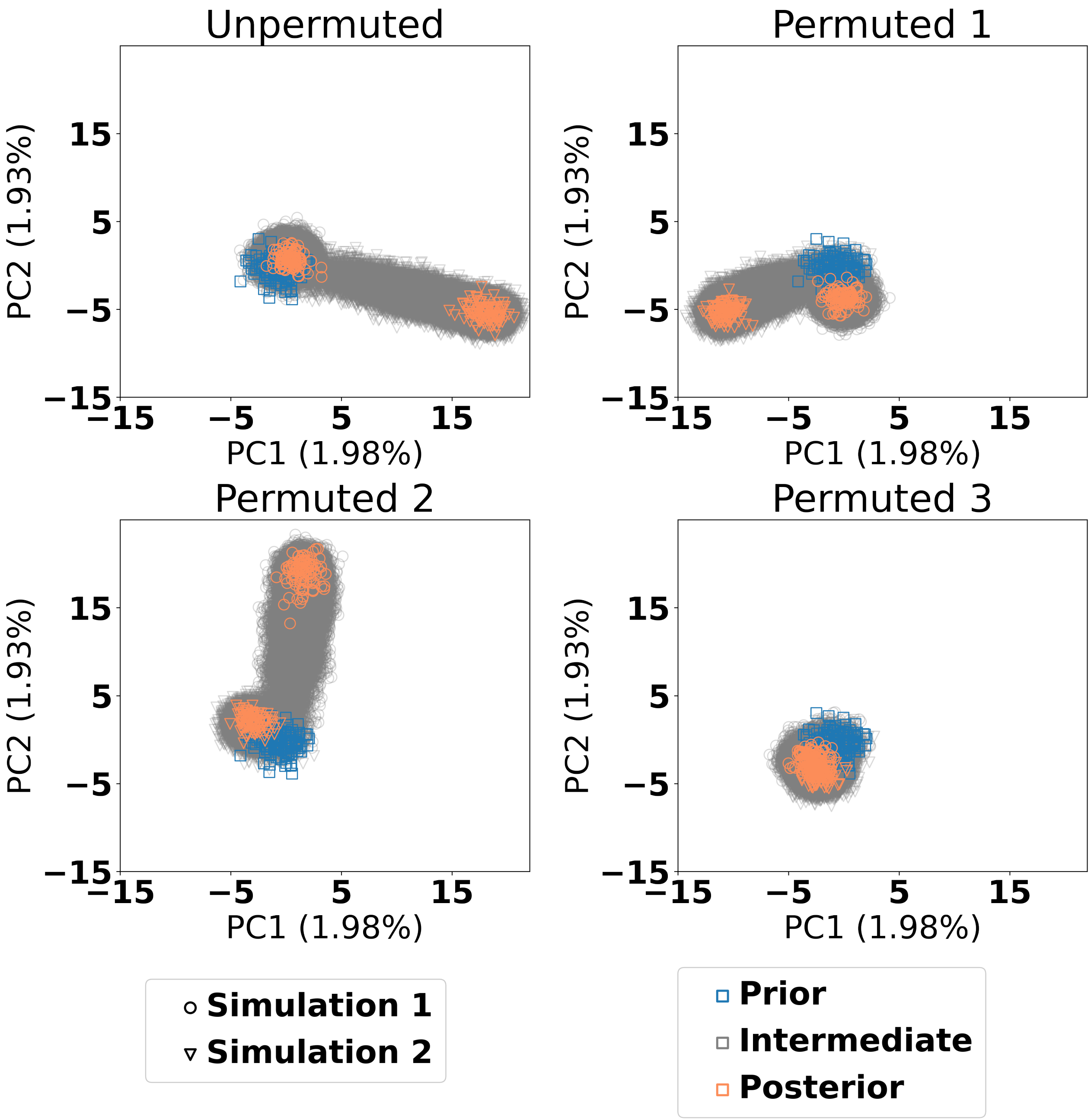


**Figure S2.** Principal Component Analysis (PCA) plot of the parameter sets used in the Bayesian calculation method where each point is a candidate parameter set. The prior points are the ones which served as a starting point for the calculation, the posterior points are the ones which had $R^2 > 0.9$, whereas all other points are intermediate points stemming from the simulations. The axes are identical for all panels and use the same ordination, making the panels directly comparable.

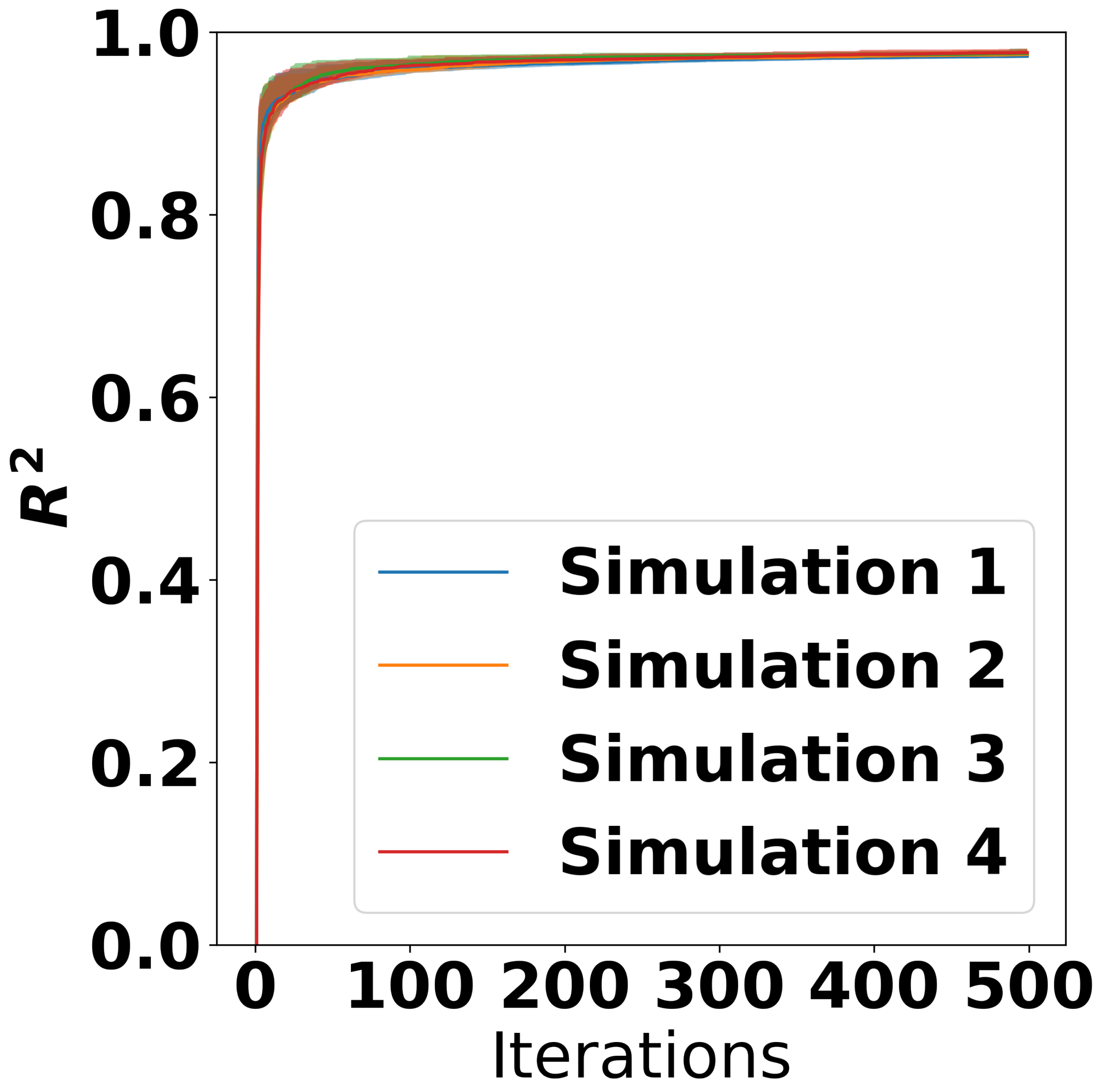


**Figure S3.** Training $R^2$ values for the Bayesian calculation method when excluding the chemostat dataset. An $R^2$ value of 1 corresponds to exact correspondence between the training data and the model predictions. The shaded regions indicate the $5^{th}$ and $95^{th}$ percentiles, whereas the solid lines indicate the median ($50^{th}$ percentile).

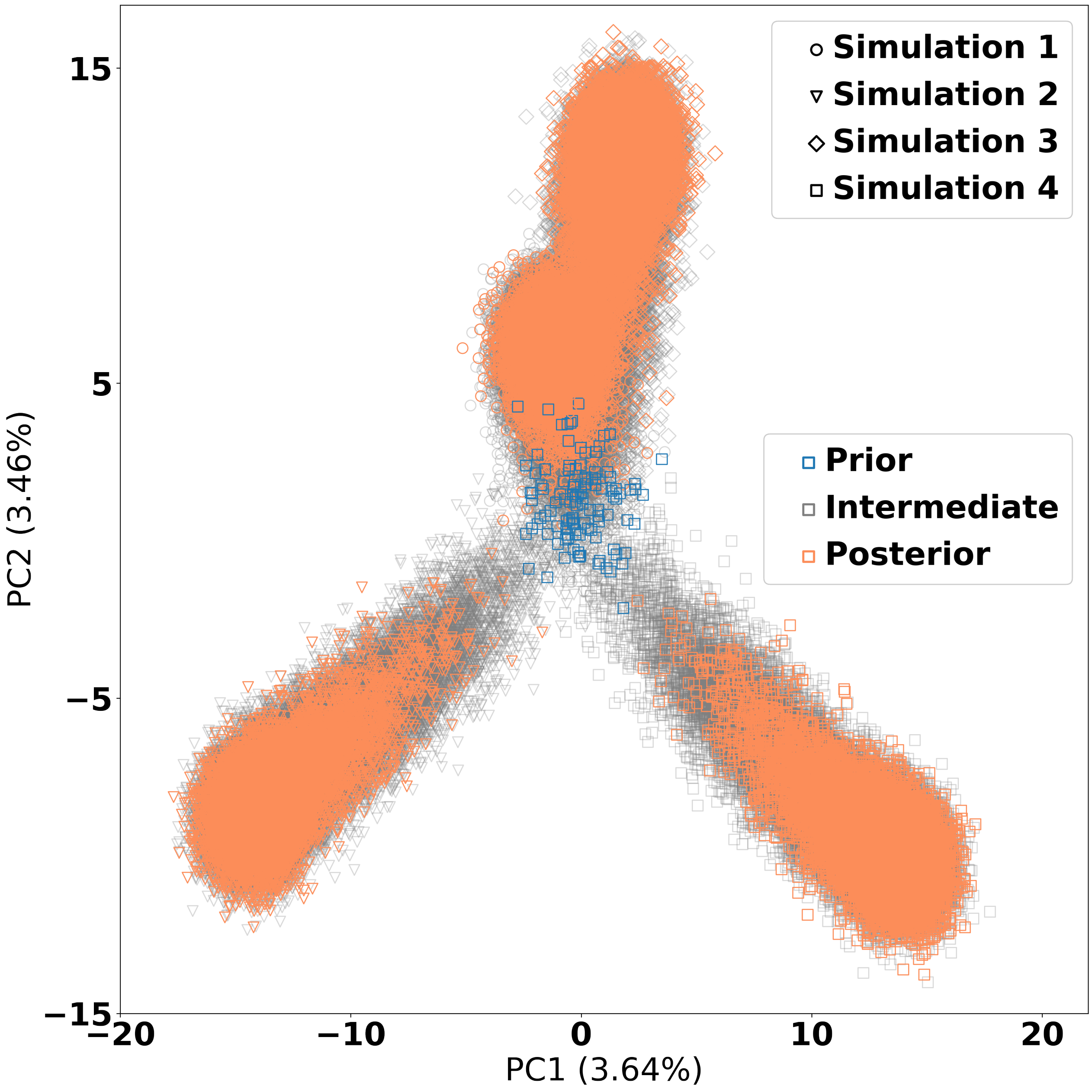


**Figure S4.** Principal Component Analysis (PCA) plot of the parameter sets used in the Bayesian calculation method when excluding the chemostat dataset. Each point is a candidate parameter set. The prior points are the ones which served as a starting point for the calculation method, the posterior points are the ones which had $R^2 > 0.9$, whereas all other points are intermediate points stemming from the simulations.

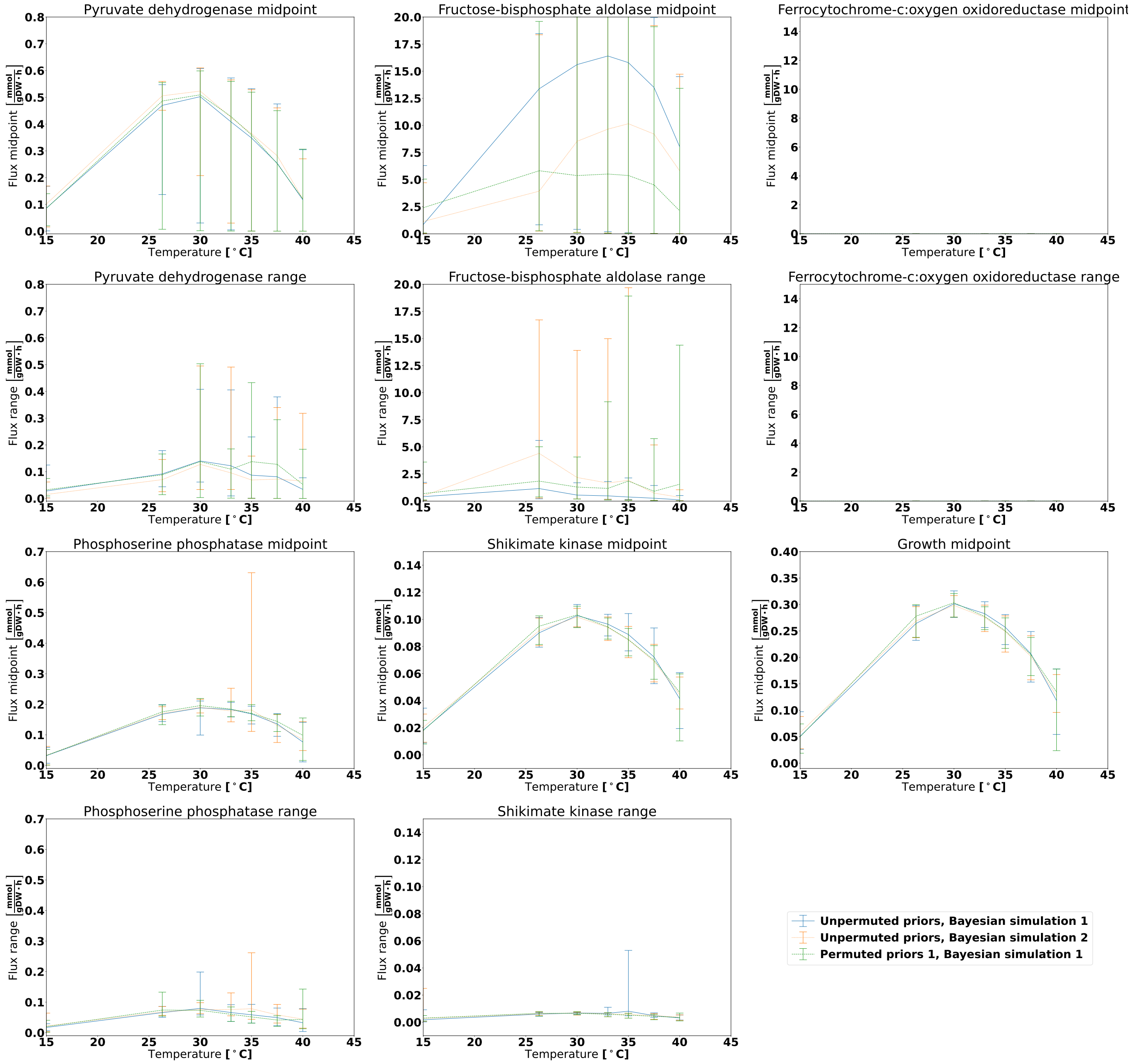


**Figure S5.** FVA analysis for the anaerobic dataset for six reactions and varying temperature when using estimated posterior distributions obtained for the Bayesian calculation method. The midpoint panels show the FVA flux midpoint, this is: The average of the maximum and minimum attainable flux given the optimization objective. The range panels show the absolute difference between the maximum and minimum flux. The lines denote the mean midpoint or range value, whereas the error bars span from the lowest to the highest observed value. The growth reaction is included for reference, and it will always display an FVA range of zero as it is the optimization target.

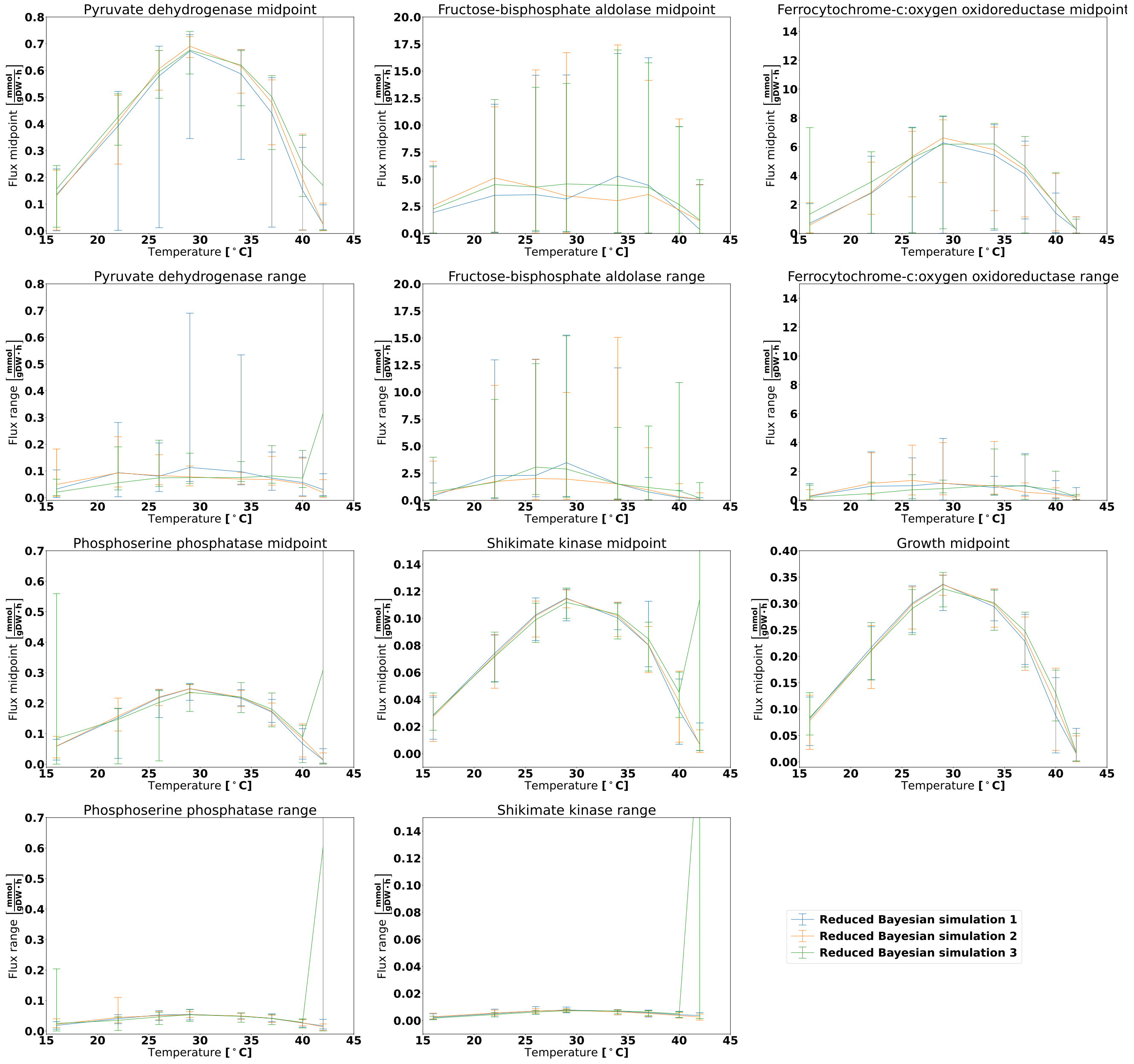


**Figure S6.** FVA analysis for the aerobic dataset for six reactions and varying temperature when using estimated posterior distributions obtained for the Bayesian calculation method without the chemostat dataset. The midpoint panels show the FVA flux midpoint, this is: The average of the maximum and minimum attainable flux given the optimization objective. The range panels show the absolute difference between the maximum and minimum flux. The lines denote the mean midpoint or range value, whereas the error bars span from the lowest to the highest observed value. The growth reaction is included for reference, and it will always display an FVA range of zero as it is the optimization target.

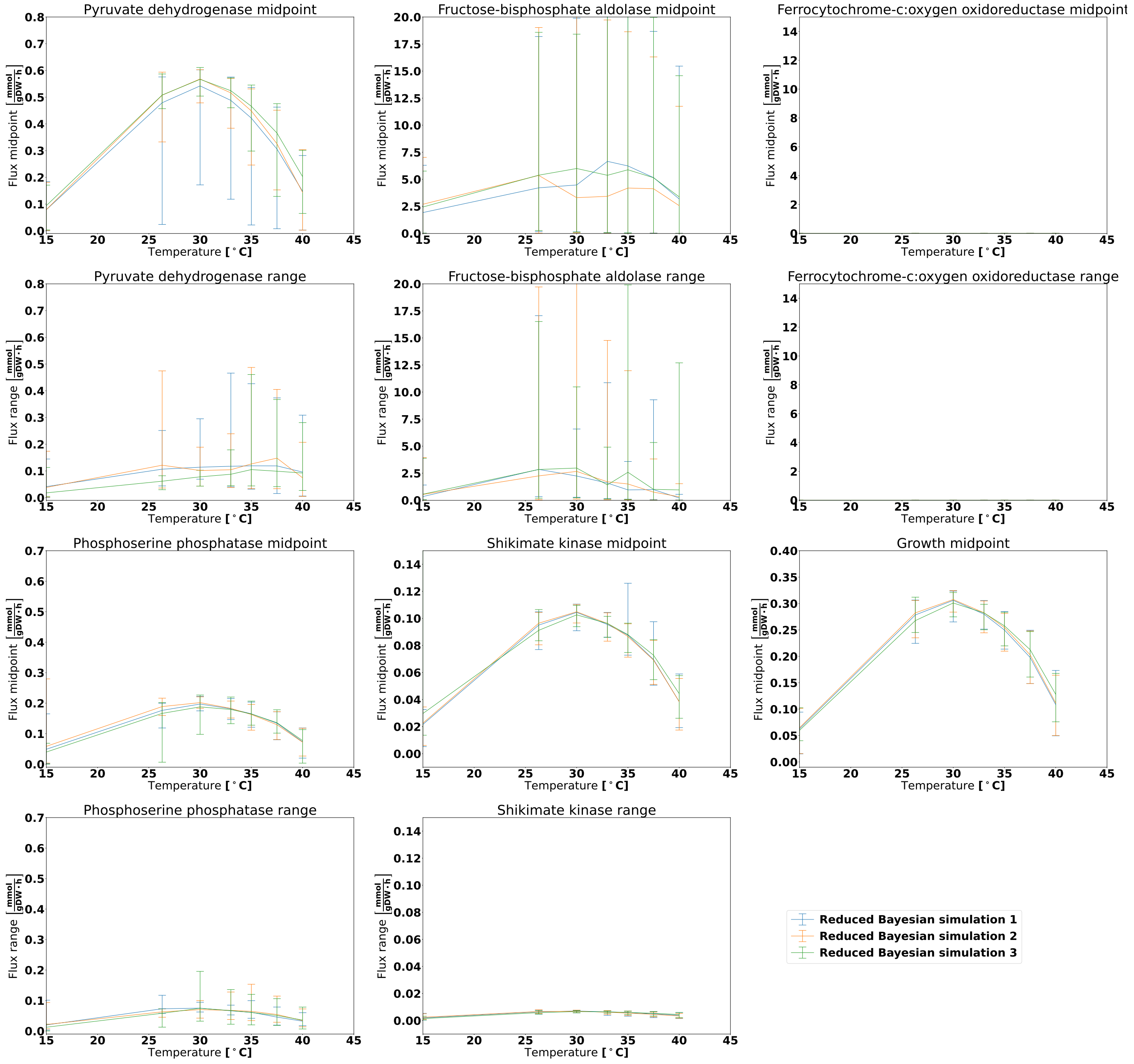


**Figure S7.** FVA analysis for the anaerobic dataset for six reactions and varying temperature when using estimated posterior distributions obtained for the Bayesian calculation method without the chemostat dataset. The midpoint panels show the FVA flux midpoint, this is: The average of the maximum and minimum attainable flux given the optimization objective. The range panels show the absolute difference between the maximum and minimum flux. The lines denote the mean midpoint or range value, whereas the error bars span from the lowest to the highest observed value. The growth reaction is included for reference, and it will always display an FVA range of zero as it is the optimization target.

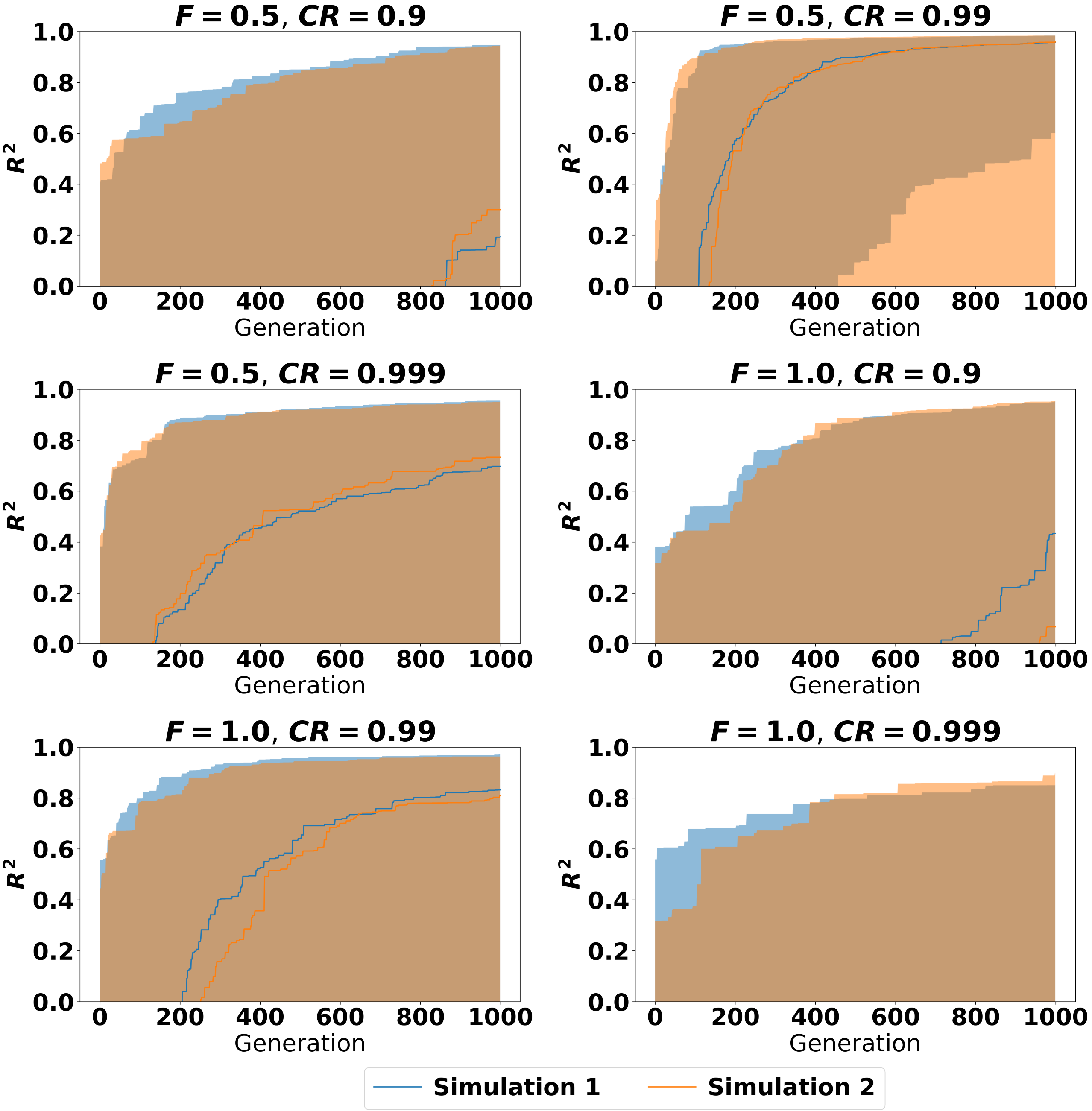


**Figure S8.** Training $R^2$ values for the evolutionary algorithm. Each panel contains the results of both simulation 1 and 2 and corresponds to a certain combination of scaling factor $F$ and crossover probability $CR$. An $R^2$ value of 1 corresponds to exact correspondence between the training data and the model predictions. The shaded regions indicate the $5^{th}$ and $95^{th}$ percentiles, whereas the solid lines indicate the median ($50^{th}$ percentile).

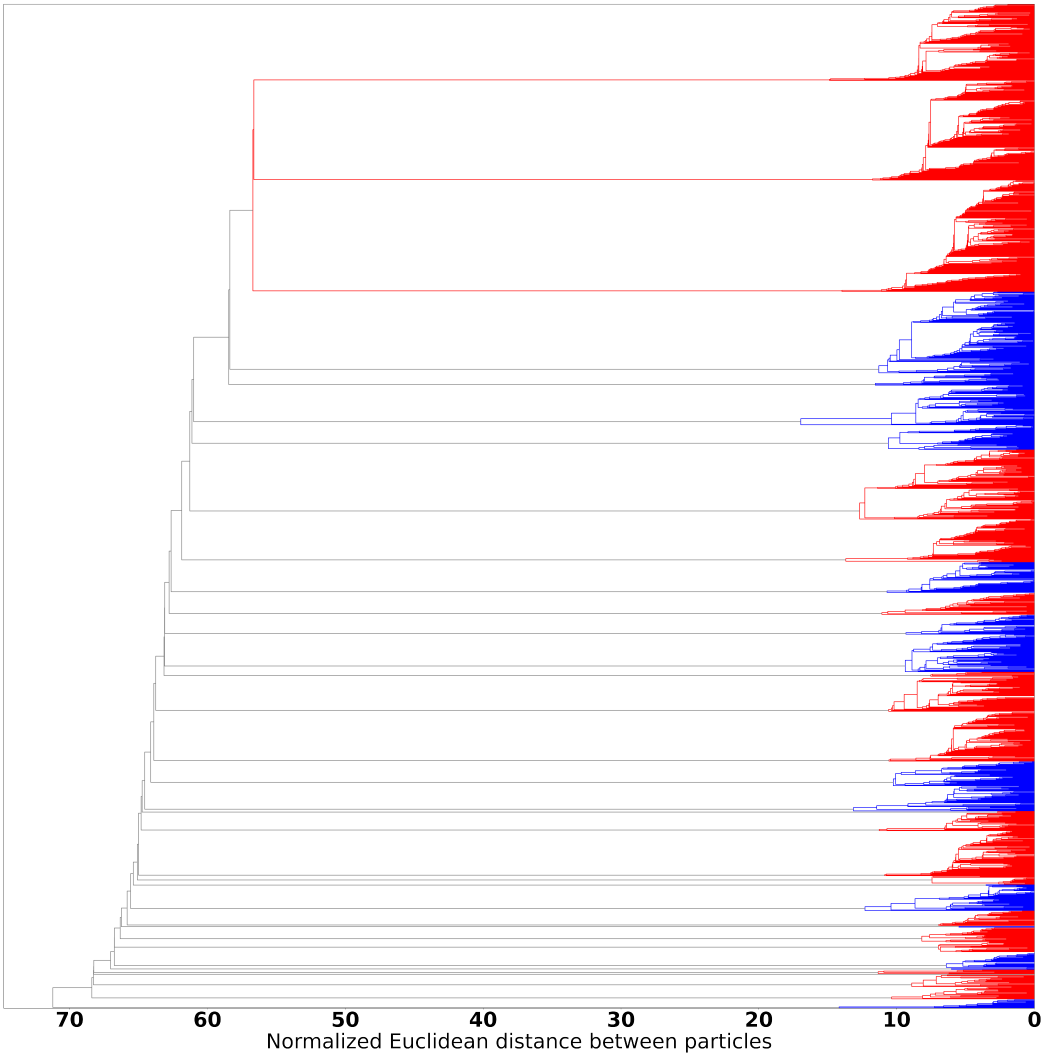


**Figure S9.** Agglomerative hierarchical clustering of the particles displayed in Figure 4 **B**. Blue branches correspond to particles from simulation 1, red branches correspond to particles from simulation 2, whereas branches which contain particles from both simulations are coloured grey.

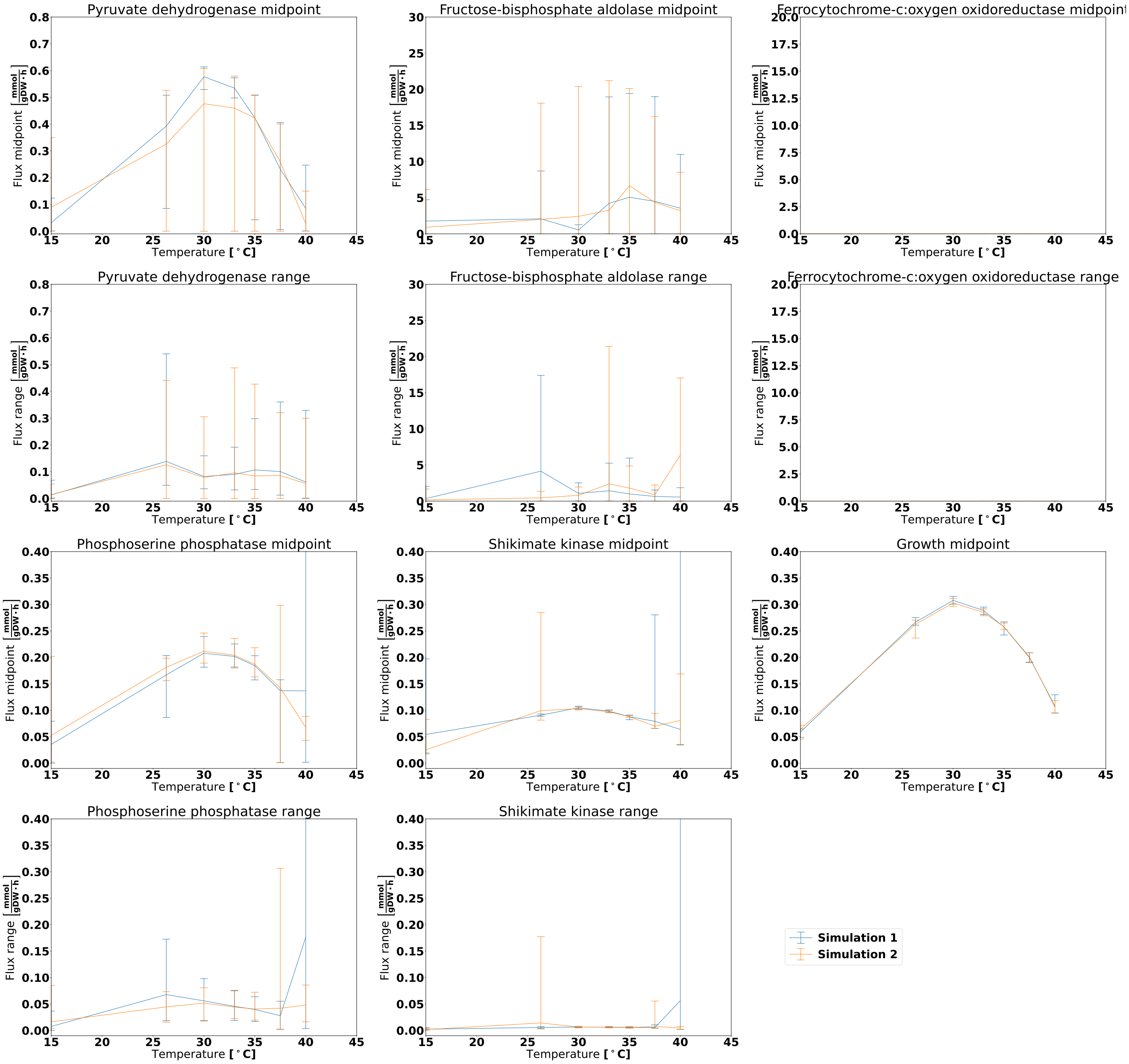


**Figure S10.** FVA analysis on the results from the evolutionary algorithm under anaerobic conditions. The particles selected for this analysis stems from the two simulations with $F = 0.5$ and $CR = 0.99$, considering only the particles with $R^2 > 0.98$. The midpoint panels show the FVA flux midpoint, ie. the average of the maximum and minimum attainable flux given the optimization objective. The range panels show the absolute difference between the maximum and minimum flux. The lines denote the mean midpoint or range value, whereas the error bars span from the lowest to the highest observed value. The growth reaction is included for reference, and it will always display an FVA range of zero as it is the optimization target.
